# A Regionally Inspired West Virginia Obesogenic Diet Induces Fat Accretion and Metabolic Dysfunction While Identifying Sex Disparity

**DOI:** 10.64898/2026.02.16.706140

**Authors:** Eric E. Kelley, Andrew P. Giromini, Brooke A. Maxwell, Amanda L. Spears, Sara E. Lewis, Sonia R. Salvatore, Marco Fazzari, Sruthi Balaji, Paolo Fagone, Emily A. Konopa, Dominique C. Saporito, Judy A. King, Francisco J. Schopfer, Nicholas K. H. Khoo, Paul McCarthy, John M. Hollander, Roberta Leonardi

## Abstract

Obesity prevalence continues to rise in the United States, with a disproportionate burden falling to West Virginia. To investigate the metabolic effects of region-specific dietary patterns, we developed the West Virginia Obesogenic Diet (WV-OD), a compositionally defined rodent diet based on nutritional analyses of meals consumed by obese individuals in the state. The WV-OD closely mirrors the macronutrient profile of the average American diet while incorporating regional features such as a greater sodium level and significantly less fiber. We compared the metabolic effects of the WV-OD to a matched control diet (WV-CD) and to a widely used high-fat diet (HFD, 60% of calories derived from fat) in male and female C57BL/6J mice. After 19 weeks, WV-OD-fed males exhibited weight gain and adiposity comparable to HFD-fed counterparts, along with glucose intolerance and hepatic triglyceride accumulation confirming the obesogenic and metabolically disruptive properties of the WV-OD. Unlike HFD-fed mice, WV-OD-fed males also displayed elevated circulating cholesterol and cholesterol esters without corresponding increases in hepatic total cholesterol. When compared to the HFD, the WV-OD did not increase uric acid or xanthine oxidoreductase (XOR) content of liver or circulation; however, both males and females on the WV-OD demonstrated trends towards elevated plasma uric acid. Interestingly, while exhibiting a similar caloric intake on either diet, the WV-OD females did not demonstrate significant fat accretion or metabolic dysfunction compared to females subjected to the 60% HFD. *In toto*, these findings: 1) establish the WV-OD as a regionally-grounded, yet broadly representative tool for modeling diet–induced obesity and metabolic dysfunction, 2) offer a physiologically relevant alternative to extreme-fat dietary models in preclinical research and 3) highlight sex-based differences in response to diet-induced obesity.

## Introduction

The global obesity crisis is rapidly becoming one of the world’s fastest-growing epidemics, with over one billion people worldwide currently living with obesity, equivalent to one in every eight individuals [1, 2]. In the United States, the prevalence of adult obesity has remained elevated and steadily increased over the past decade, rising from 37.7% in 2013–2014 to 40.3% in 2021–2023 [3]. When combined, the clinical classifications of “overweight” and “obese”, identify about three in four adults [4] and, based on current trends, projections suggest that by 2050, one in three adolescents and two in three adults over the age of 25 will be overtly obese in the United States [5]. The impact of these trends is manifested to the greatest extent in the state of West Virginia in which 41.4% of the population is considered obese.

To investigate the pathophysiology of obesity, the use of rodent models fed high-fat diets (HFDs), typically ranging from 40% to 60% kcal from fat, has become standard [6–8]. In 1999, scientists at Research Diets formulated a purified diet containing 349 g of fat per kilogram, primarily from lard, corresponding to 60% of total caloric intake. This diet was designed to accelerate the onset and severity of obesogenic phenotypes compared to a 45% kcal fat diet. Male and female mice on the 60% HFD become obese more rapidly, allowing for shorter study durations and contributing to the diet’s widespread adoption [8–11]. However, the extreme fat content and reliance on a single fat source do not reflect typical human dietary patterns. According to the 2021 National Health and Nutrition Examination Survey (NHANES), the average American diet consists of approximately 49% carbohydrates, 35% fat, and 16% protein, with fats sourced from a mix of animal and vegetable products.

Initial efforts to simulate human obesogenic eating patterns relied on cafeteria diets, in which rodents self-selected from a variety of highly palatable processed foods. While these models effectively induce hyperphagia and rapid weight gain, regional differences in the nutritional composition of similar food items and uncontrolled intake patterns limit their reproducibility across studies [12–14].

To better mimic human dietary habits and improve experimental consistency, several purified Western-style diets have been developed in recent years. The Total Western Diet (TWD), formulated using NHANES median nutrient intakes, incorporated multiple fat sources and an adjusted micronutrient mix [15]. However, testing the TWD in male and female mice did not induce robust obesity or glucose intolerance in either sex [16]. The absence of a strong metabolic phenotype was likely due to suppressed hyperphagia, possibly linked to the modified micronutrient profile. The Standard American Diet (SAD) was later developed to include various fat sources, including trans-fats and omega-6 polyunsaturated fatty acids (PUFAs), to promote a pro-inflammatory nutrient profile [17]. The SAD consistently led to increased body weight, fat mass accumulation, and impaired glucose tolerance in both male and female mice compared to standard chow-fed controls [17, 18]. Following the FDA’s trans-fat ban, which took effect in 2018, a modified version of the SAD, referred to as mSAD, was developed [19]. The mSAD replaced trans fats with palm oil, substituted sucrose with high-fructose corn syrup, reduced total fiber content, and increased the proportion of calories from fat to 49.2%, compared to 35.6% in the original SAD. This formulation produced rapid hyperglycemia and insulin resistance in male mice within just four weeks of feeding and led to robust weight gain, compared the standard rodent diet AIN-93M which, though not ingredient-matched, is commonly used as a reference diet in rodent studies [20].

While these diets more closely approximate human dietary patterns in mice, they rely on national dietary averages and do not account for regional variation in food intake or obesity burden. West Virginia currently has the highest prevalence of adult obesity in the United States (∼41.4%) and one of the greatest prevalences of adult diabetes (∼15%) [21, 22].

To better model the dietary drivers of obesity in this high-risk population, we developed the West Virginia Obesogenic Diet (WV-OD) based on clinical dietary surveys from overweight patients in our region. The WV-OD is a compositionally defined diet containing 39% kcal from multiple fat sources (lard, butter, and soybean oil), 47% kcal from carbohydrates (22% from sucrose), and 14% kcal from protein. Compared to a matched control diet (WV-CD), the WV-OD also contains lower fiber and higher sodium. In this study, we tested the WV-OD in both male and female mice, evaluating its effects relative to the WV-CD and to the benchmark 60% HFD. We report that the WV-OD induced a robust male-specific obese phenotype comparable to that observed in the 60% HFD group. This phenotype was associated with glucose intolerance, albeit to a lesser extent than with the 60% HFD, and with plasma cholesterol levels exceeding those in the 60% HFD group. This regionally-grounded model offers a new tool for studying how Western dietary patterns contribute to obesity and metabolic dysfunction in high-risk populations.

## Materials and Methods

### Reagents

Uric acid (UA), xanthine, allopurinol, and oxonic acid were purchased from Sigma-Aldrich. β-Nicotinamide adenine dinucleotide was purchased from MP Biochemicals. All other reagents were of analytical grade or better and were purchased from Fisher Scientific or Sigma-Aldrich.

### Diet analysis and formulation of the WV-OD

Samples of daily meals (8 breakfasts, 10 lunches, and 9 dinners) consumed by local overweight (BMI > 25) individuals were gathered during routine nutritional counseling sessions with a dietitian nutritionist (**Supplementary Table S1**). All information was fully de-identified before being shared with the research team, and no personal data were accessed or recorded. The initial meal analysis focused on determining the proportion of calories derived from total fat, saturated fat, protein, total carbohydrates and sugars, as well as the amount of sodium, fiber and cholesterol present in food items (**Fig. 1A**). This assessment relied on available food labels for exact or comparable items. In instances where specific details were unavailable, we made certain assumptions: serving sizes were set at twice the indicated amount on the food label, and meat and egg preparation was assumed to involve 2 tablespoons of unsalted butter. For each meal, we also calculated the percentage of total fat sourced from meat plus eggs, dairy, and vegetable oil (**Fig. 1B**). This information guided our selection and proportions of fat sources included in the WV-OD and control diet (WV-CD), with lard used to approximate the fat supplied by meat and eggs (40% of total fat), butter used to represent fat from dairy (30% of total fat), and corn oil (30% of total fat) used to approximate all vegetable oil. Sugar-sweetened beverages (SBB) were inconsistently annotated during the interviews; however, considering the prevalent consumption of sugar-sweetened beverages reported by West Virginia residents [23], we created a “model” SSB by averaging the nutritional composition of all reported SSBs (**Supplementary Table S1**). We then calculated the percentage of calories attributed to different macronutrients using two distinct calculation methods (**Supplementary Fig. S1**). The first method combined the nutritional content of 2 SSBs with the average nutritional composition of breakfasts, lunches, and dinners, reflecting a three-meal-per-day consumption pattern. This approach was also used to estimate the daily intake of fiber, cholesterol and sodium (**Table 1**). The second method entailed adding 1 SSB separately to each lunch and dinner, recalculating their nutritional compositions, and subsequently averaging the nutritional contents of all breakfast, lunch (with an added SSB), and dinner (also with an included SSB) meals together. This approach determined the percentage of calories derived from total carbohydrates, sugars, proteins, total fat and saturated fat in an average meal without relying on a fixed assumption of a three-meal per day pattern. Indeed, some individuals who provided the meal information reported having only had two meals in a day. The results obtained with the two methods were very similar, with differences ≤ 1% (**Supplementary Fig. S1**). Nutrient contribution to calorie intake, as calculated by the second method, and the daily intake of sodium, cholesterol and fiber calculated with the first method were compared to the Recommended Daily Allowances (RDAs) for the US population [24] (**Table 1**). The formulation of the WV-OD and the WV-CD was guided by the differences observed between the values obtained from the sample meals and the RDAs. Additionally, we incorporated nutritional recommendations for mice wherever available [25]. The WV-CD and WV-OD formulations adhered to recommended mineral and vitamin doses for mice, matching, with the exception of sodium, amounts contained in Research Diets D12492 and D12450J. Since specific recommendations for daily fiber intake in this species are unavailable, we matched the total fiber content (cellulose plus inulin) in the WV-OD to levels commonly used in other rodent diet formulations and increased the amount in the WV-CD threefold. The WV-OD and WV-CD were produced by Research Diets with numbers D23031308 and D23031309, respectively.

**Figure 1.**
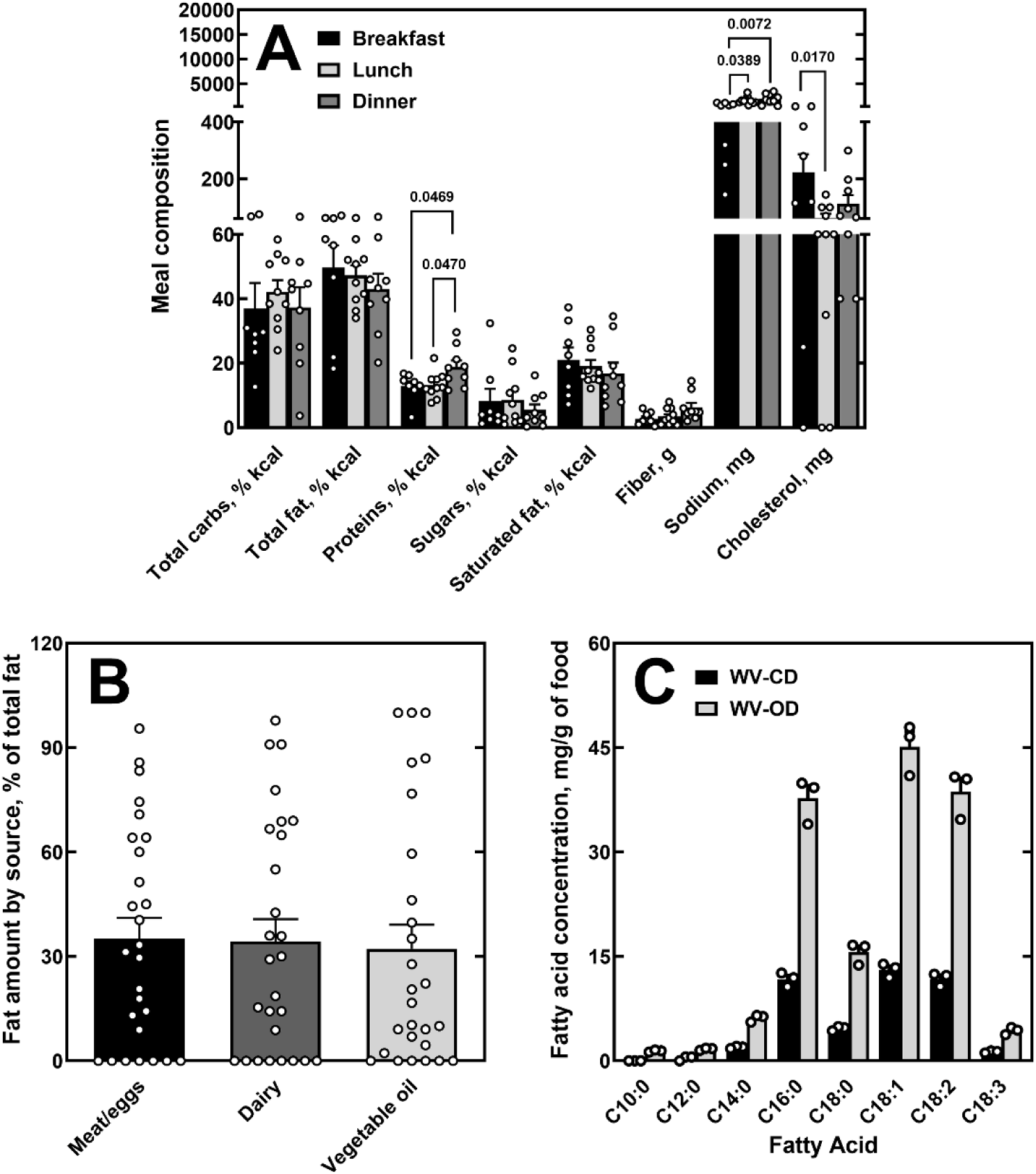
Nutritional analysis and fatty acid composition of the WV diets. (**A**) Macronutrient composition of 27 patient-reported meals, separated by meal type. (**B**) Contribution of different fat sources to total dietary fat in the analyzed meals. (**C**) Fatty acid composition of the WV-OD and WV-CD based on GC-FID analysis. Data shown as mean ± SEM. Individual data points represent independent meals (A and B) or diet batches (C).

**Table 1.** Summary of the average nutritional composition of the 27 meals analyzed + 2 SSB, with the USDA recommended values and the mean values found in the American diet as reference.

| Nutrients | Recommended values<sup>#</sup> | Mean values in the American diet (2021-2023)<sup>##</sup> | Mean values in the analyzed meals |
| --- | --- | --- | --- |
| Total carbohydrates, % kcal | 45-65 | 45 | 49 |
| Total fat, % kcal | 20-35 | 37 | 39 |
| Protein, % kcal | 10-35 | 16 | 14 |
| Sugars, % kcal | <10<sup>a</sup> | 19<sup>\$</sup> | 22 |
| Saturated fat, % kcal | <10 | 12 | 15 |
| Cholesterol, mg/day | N/A<sup>b</sup> | 321 | 400 |
| Sodium, mg/day | 2300 | 3212 | 4315 |
| Fiber, g/day | 22-34 | 16 | 13 |

<sup>#</sup> Age group >20, males and females, Dietary Guidelines for Americans, 2020-2025, https://www.dietaryguidelines.gov
<sup>##</sup> What We Eat in America, NHANES 2021-2023, https://www.ars.usda.gov/northeast-area/beltsville-md-bhnrc/beltsville-human-nutrition-research-center/food-surveys-research-group/docs/wweia-data-tables/
<sup>\$</sup>Value was recalculated as % of kcal consumed for males and females, 20 and over in age
<sup>a</sup>Value specifically refers to added sugars
<sup>b</sup>The new guidelines suggest keeping cholesterol consumption “as low as possible without compromising the nutritional adequacy of the diet”. Dietary Guidelines for Americans, 2020-2025, https://www.dietaryguidelines.gov

### Mouse studies

All animal studies were conducted under the approval of the West Virginia University Institutional Animal Care and Use Committee (protocol# 1604002026). Mice were housed in an animal facility kept at a room temperature of 22.0 ± 0.2 °C, room humidity of 40% ± 2%, and a 12-h light/12-h dark cycle, with the dark cycle starting at 18:00 h. Male and female C57BL/6J mice were fed a 60% HFD (Research Diets D12492, referred to as HFD) or the WV-OD for 19 weeks, beginning at 6 weeks of age. Control mice were fed matched control diets containing 10% of calories from fat (Research Diet D12450J, referred to as CD) or the WV-CD, which contains 15% of calories from fat. Data from mice fed the 60% high-fat diet (HFD) and its corresponding control diet (CD) were previously reported in [29]. The present study includes all mice from the previously reported HFD and CD cohorts that completed 18 weeks on their respective diets (all females and a subset of males). In the previous study, these data were analyzed to compare responses to the HFD between males and females, whereas in the present study they were reanalyzed to compare the WV-OD to the HFD and their respective control diets within each sex. The WV-OD and WV-CD data are entirely new, and the corresponding cohorts were studied alongside the HFD and CD groups, with overlapping experimental periods. Diets and water were supplied *ad libitum* for the entire study. Mouse weights and food consumption were recorded weekly. Lean and fat mass were assessed by EchoMRI after 16 weeks of diet intervention. For glucose tolerance tests (GTTs), mice were fasted for 7 h starting at 7:00 A.M. and then injected intraperitoneally with glucose (1.3 g/kg). This dose remained constant throughout the study. Blood glucose was measured before (t=0) and at 20, 40, 60, and 120 min after glucose injection using an Accu-Chek Guide Me glucometer. For tissue harvest, the mice were fasted for 7 hours and anesthetized with isoflurane, followed by blood collection via cardiac puncture and subsequent removal of the organs of interest. Tissue samples were quickly weighed before flash-freezing in liquid nitrogen, then stored at -80 °C until analysis. Blood was collected using heparinized syringes and centrifuged at 8,600 × g for 6 minutes at 4 °C to isolate plasma.

### Liver histology and measurement of xanthine oxidoreductase activity

Livers were harvested as described above and immediately fixed in 10% neutral-buffered formalin for one week at 4 °C. Fixed tissues were then embedded in paraffin and sectioned at 5 μm thickness. Hematoxylin and eosin (H&E) staining was performed according to standard protocols by the WVU Histopathology Core Facility. Images were acquired using an Olympus BX53 light microscope with CellSens imaging software. Macrovesicular steatosis was semi-quantitatively scored as previously described [26].

Xanthine oxidase (XO) activity in plasma and xanthine oxidoreductase (XOR, XO and xanthine dehydrogenase (XDH)) activity in liver tissue were measured as we previously described [27–29].

### Lipid measurements in plasma and liver

Plasma concentrations of triglycerides, cholesterol, and cholesterol esters were determined by LC-MS/MS, as previously described [29]. Liver triglycerides were extracted and quantified using the protocol reported by Shumar et al. [30], while total cholesterol was extracted using the same protocol but quantified using a commercial kit (Thermo Scientific) according to the manufacturer’s instructions.

### Fatty acid analysis of the WV-OD and WV-CD diets

Lipids were extracted from 3 independent batches of WV-OD and WV-CD diets, purchased within 4 months of each other, using a modified version of a previously described protocol [30]. Briefly, ∼50 mg of food were crushed into small pieces and incubated overnight at room temperature in a 2.4:1 (v:v) mixture of methanol containing 2% acetic acid and chloroform. To induce phase separation, 1.2 ml of water and 1.5 ml of chloroform were added to the extract, followed by centrifugation. The lower (organic) phase was collected, and the upper (aqueous) phase was re-extracted with 1 ml of chloroform. All chloroform extracts were pooled and dried under a stream of nitrogen. Fatty acid methyl esters (FAMEs) were prepared as described by Lepage et al. [31]. Dried lipid extracts were resuspended in 3 ml of 2:1 methanol:hexane (v:v), and 0.5 ml aliquots were mixed with 100 µg of tridecanoic acid (internal standard) and 100 µl of 1 M acetyl chloride in dichloromethane. Samples were heated at 100 °C for 1 h, cooled to room temperature, and extracted with 1 ml of hexane and 2 ml of 6% potassium carbonate solution. The upper (hexane) phase containing the FAMEs was collected for analysis. FAMEs were analyzed by gas chromatography with flame ionization detection (GC/FID) using an Agilent 7890 GC system equipped with a DB-23 capillary column (60 m × 0.25 mm). Injections were performed in split mode using helium as the carrier gas at a flow rate of 2.0 ml/min. Oven conditions were as follows: initial isotherm at 60 °C for 1 min; ramp at 25 °C/min to 185 °C; ramp at 1 °C/min to 215 °C; ramp at 25 °C/min to 240 °C; final hold at 240 °C for 3 min. The injector and detector were maintained at 250 °C and 280 °C, respectively. FAMEs were identified by comparing retention times to a commercial FAME standard mix (Supelco), and quantification was performed by normalizing peak areas to the known amount of internal standard and the initial diet weight.

### Statistical analysis

All statistical analyses were performed using Prism 10.2.3 (GraphPad, San Diego, CA). Data are expressed as the mean ± SEM. Comparisons were conducted using one-way or two-way ANOVA, as indicated, followed by Tukey’s multiple comparisons test, or by linear regression with selected slope comparisons. Differences between groups were considered significant at p < 0.05. For regression-based comparisons, slopes also had to differ by at least 10% to be considered biologically meaningful. Outliers were identified using the ROUT method with a false discovery rate (Q) of 1%. No data exclusion criteria were pre-established.

## Results

### Formulation of the WV-OD and effect on weight gain and body composition

To develop a regionally grounded model of dietary patterns in high-risk populations, we first analyzed the macronutrient composition of 27 representative meals consumed by overweight individuals in West Virginia. The composition of the 27 meals analyzed, excluding sugar-sweetened beverages (SSB), is shown in **Fig. 1A**. Notably, the percentage of calories derived from total carbohydrates, simple sugars, and fat was comparable across all meal types. Similarly, the proportion of saturated fat and the amount of fiber remained consistent. Conversely, differences were found in the percentage of calories derived from proteins, which was significantly greater in dinner meals, and breakfast meals contain lesser sodium levels but greater cholesterol amounts, compared to lunch and dinner. Frequent consumption of sugar-sweetened beverages (SSBs) is linked to an increased risk of obesity and metabolic syndrome [32–34]. Considering the high prevalence of daily SSB consumption among West Virginia adult residents [23, 35], we included two SSBs in the meal composition analysis. As shown in **Table 1**, the percentage of calories derived from total carbohydrates, sugars, total fat, saturated fat, and protein in the 27 patient meals closely aligned with the most recent composition data for the average American diet, based on the 2021-2023 NHNES. Both the average American diet and the meals we analyzed greatly exceeded recommended levels for calories from sugars and contained greater levels of saturated fat. Daily sodium intake surpassed current recommendations, and daily cholesterol intake surpassed the former guideline of 300 mg/day, which has been replaced with advice to keep cholesterol intake “as low as possible without compromising the nutritional adequacy of the diet” [24]. Fiber intake fell below recommended levels in both the average American diet and the meals we analyzed, with the fiber in the latter trending even less. Based on the meal analysis, we designed a rodent diet, the West Virginia Obesogenic Diet (WV-OD), with 39% of its calories derived from fat and 22% of calories derived from sucrose (**Table 2**). As a source of fat, we selected lard, butter, and corn oil in a rounded 40:30:30 ratio to mirror the average fat composition observed in the analyzed meals (**Fig. 1B**). A high ω-6/ω-3 ratio has long been associated with the pro-inflammatory nature of Western eating patterns [36], although more recent studies suggest that this ratio alone may not fully capture inflammatory risk [37]. As a reference, the mean ratio of 18:2 (predominantly linoleic acid, an ω-6 fatty acid) to 18:3 (predominantly linolenic acid, an ω-3 fatty acid) in the WV-OD was 9.0 (**Fig. 1C**), closely aligning with the estimated 9.7 ratio in the average American diet. This similarity supports the notion that the WV-OD, while regionally informed, mirrors key macronutrient and fatty acid characteristics of the broader U.S. diet and is not compositionally idiosyncratic.

**Table 2.** Composition of the WV-OD, WV-CD, HFD, and CD.

| **Macronutrients** | **WV-OD**<br>**(D23031308)** |  | **WV-CD**<br>**(D23031309)** |  | **HFD**<br>**(D12492)** |  | **CD**<br>**(D12450J)** |  |
| --- | --- | --- | --- | --- | --- | --- | --- | --- |
|  | g | kcal<br>(% of total) | g | kcal<br>(% of total) | g | kcal<br>(% of total) | g | kcal<br>(% of total) |
| Corn starch | 117 | 11.7 | 498 | 49.8 | 0 | 0 | 506 | 49.9 |
| Sucrose | 210 | 22.0<sup>a</sup> | 60 | 7.0<sup>a</sup> | 68.8 | 7.8<sup>a</sup> | 68.8 | 7.8<sup>a</sup> |
| Maltodextrin 10 | 125 | 12.5 | 125 | 12.5 | 125 | 12.3 | 125 | 12.3 |
| Cellulose | 33 | 0 | 100 | 0 | 50 | 0 | 50 | 0 |
| Inulin | 17 | 0.7 | 50 | 1.9 | 0 | 0 | 0 | 0 |
| *Total carbohydrates* | *502* | *47* | *833* | *71* | *244* | *20* | *750* | *70* |
| Casein | 138 | 13.8 | 138 | 13.8 | 200 | 19.7 | 200 | 19.7 |
| L-Cystine | 2 | 0.2 | 2 | 0.2 | 3 | 0.3 | 3 | 0.3 |
| *Protein* | *140* | *14* | *140* | *14* | *203* | *20* | *203* | *20* |
| Soybean oil | 51 | 11.5 | 19.5 | 4.4 | 25 | 5.5 | 25 | 5.5 |
| Lard | 72 | 16.2 | 27 | 6.0 | 245 | 54.3 | 20 | 4.4 |
| Butter | 51 | 11.5 | 19.5 | 4.4 | 0 | 0 | 0 | 0 |
| *Total fat<sup>b</sup>* | *174* | *39* | *66* | *15* | *270* | *60* | *45* | *10* |
| **Micronutrients** |  |  |  |  |  |  |  |  |
| Mineral mix<br>S10026/S10026A<sup>c</sup> | 5 | 0 | 5 | 0 | 10 | 0 | 10 | 0 |
| Sodium chloride | 4.4 | 0 | 2.6 | 0 | 0 | 0 | 0 | 0 |
| Calcium phosphate | 13 | 0 | 13 | 0 | 13 | 0 | 13 | 0 |
| Calcium carbonate | 5.5 | 0 | 5.5 | 0 | 5.5 | 0 | 5.5 | 0 |
| Potassium citrate<br>monohydrate | 16.5 | 0 | 16.5 | 0 | 16.5 | 0 | 16.5 | 0 |
| Vitamin mix<br>V10001 | 10 | 1 | 10 | 1 | 10 | 1 | 10 | 1 |
| Choline bitartrate | 2 | 0 | 2 | 0 | 2 | 0 | 2 | 0 |
| **Energy density**<br>**(kcal/g)** |  | 4.6 |  | 3.7 |  | 5.2 |  | 3.8 |

<sup>a</sup>This value includes the amount of sucrose present in the vitamin mix.
<sup>b</sup>Butter and lard contain approximately 0.22 and 0.07 g of cholesterol per 100 g, respectively. Additionally,
casein also contains 0.02 g of cholesterol per 100 g. Based on these estimates, the WV-OD and WV-CD
contain 0.022% and 0.008% cholesterol, respectively, whereas the HFD and CD contain 0.027% and 0.005%.
<sup>c</sup>Diets WV-OD and WV-CD were formulated using mineral mix S10026A.

The control diet (WV-CD) for the WV-OD was formulated with 15% of total calories from fat, using the same fat sources in the same 40:30:30 ratio. It also included 7% of total calories from sucrose, reflecting the average contribution of simple sugars across the 27 meals analyzed prior to inclusion of sugar-sweetened beverages (SSBs) (**Fig. 1A**). Sodium content was maintained at 1 g/kg in the CD, consistent with standard mouse diets. In contrast, sodium was increased to 2 g/kg in the WV-OD to model the approximately twofold greater daily sodium intake extrapolated from the analyzed meals relative to recommended intake levels (**Table 1**). Total fiber content in the WV-OD was set at 50 g, which is typical of purified rodent diets, including both HFD and CD formulations. However, the fiber source consisted of a 2:1 mixture of cellulose to inulin, a soluble fiber shown to decrease loss of intestinal mass, compared to cellulose alone [38, 39]. To reflect the gap between the recommended fiber intake for adult Americans and the lower estimated fiber intake derived from the analyzed meals (**Table 1**), fiber content in the CD was increased to 150 g while maintaining the same insoluble-to-soluble fiber ratio as in the WV-OD.

The effect of feeding the WV-OD on weight gain and body composition was tested in male and female C57Bl/6J mice and compared to the WV-CD. Parallel cohorts of mice were fed the HFD and its matched CD. Over 19 weeks, male mice fed the WV-OD gained significantly more weight than those fed the WV-CD, confirming the obesogenic nature of the WV-OD (**Fig. 2A**). Notably, the rate of weight gain in the WV-OD group was comparable to that observed in mice fed the 60% HFD and correlated with a similar accumulation of body fat (**Fig. 2C**). Consistent with this finding, energy intake was similar between the WV-OD and HFD groups (**Fig. 2B**), owing to a greater food consumption in the WV-OD-fed mice (17.3 g/mouse/week vs. 14.7 g/mouse/week, p < 0.0001, **Supplementary Fig. S2A**), which compensated for the WV-OD lower energy density (4.6 kcal/g; **Table 2**) relative to the 60% HFD (5.4 kcal/g). Overall, while mice fed the WV-OD and WV-CD consumed similar amounts of food by weight, food intake in the HFD-fed group was less than in mice fed the matched CD, a previously documented effect likely attributable to the greater difference in fat content between those diets [40].

**Figure 2.**
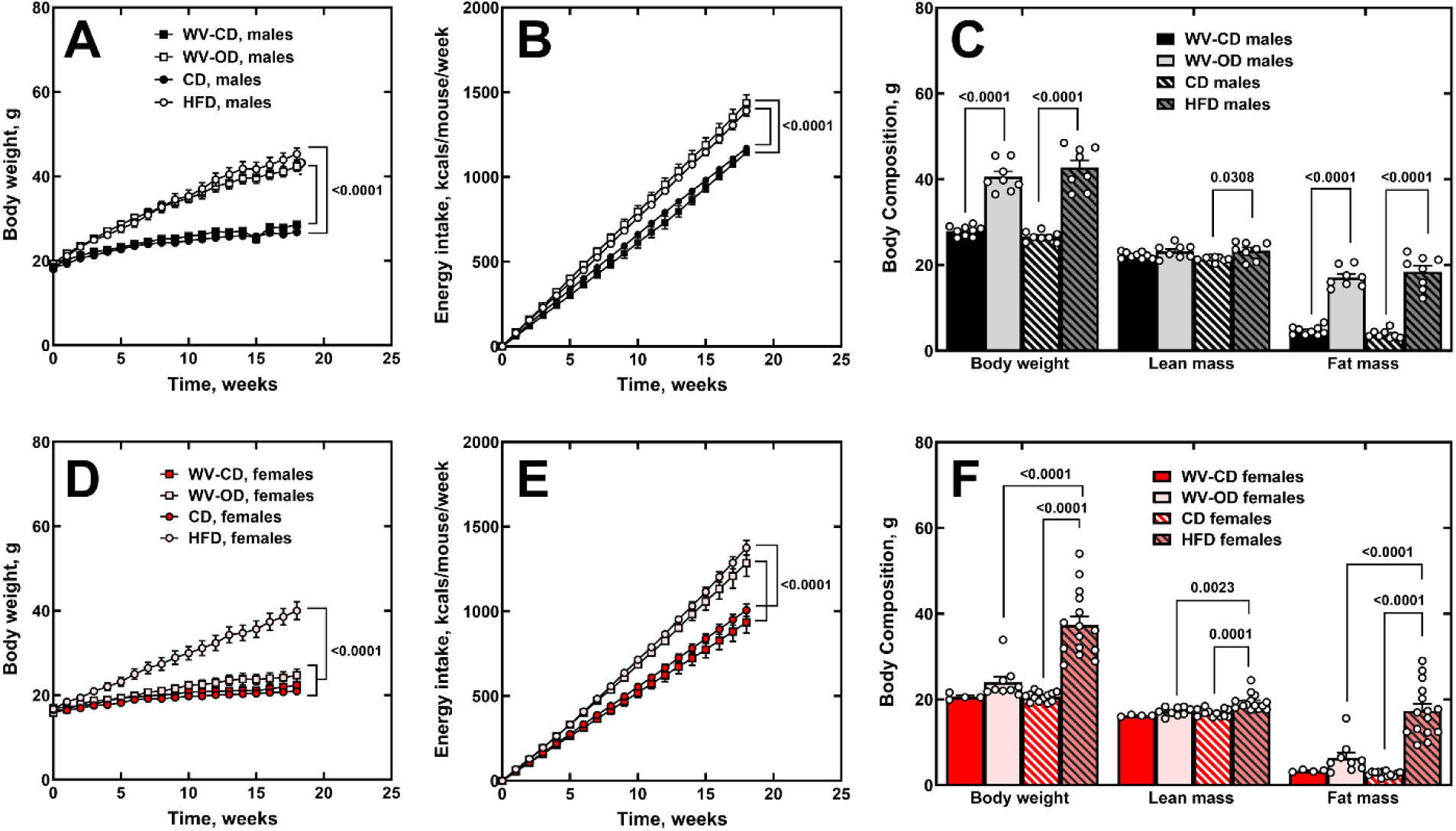
WV-OD induces robust weight gain and adiposity in male mice. (**A**, **D**) Body weight gain and (**B**, **E**) cumulative energy intake over the first 18 weeks of diet feeding in (**A**, **B**) male and (**D**, **E**) female mice. (**C**, **F**) Body composition analysis in (**C**) male and (**F**) female mice after 16 weeks on the diet. Mice were harvested at week 19. Data are shown as the mean (bars) of individual animals (circles) ± SEM. (A, D) Two-way ANOVA; (B, E) linear regression with slope comparison; (C, F) one-way ANOVA. *p*-values are shown for statistically significant comparisons.

In contrast to males, WV-OD-fed female mice trended towards greater weight gain, yet, did not gain statistically significant more weight than their WV-CD-fed counterparts, despite a substantially greater energy intake (71.7 kcal/mouse/week vs. 51.6 kcal/mouse/week, p < 0.0001) (**Fig. 2D**, **E**). Notably, this energy intake was comparable to that of females fed the HFD, who did exhibit significant weight gain and fat mass accumulation compared to all other groups (**Fig. 2D**, **F**). While these data demonstrated that the WV-OD promotes weight gain in a male-specific manner and that this gain was similar to the 60% HFD, it also shows an amplification of the resistance/time lag for fat accretion/metabolic dysfunction characteristic of female mice in general, and C57BL/6J mice in particular.

### Effect of the WV-OD on glucose tolerance and lipids in the liver and plasma

As expected, mice of both sexes fed the HFD for 10 weeks exhibited glucose intolerance and had significantly greater fasting blood glucose levels compared to CD-fed controls (**Fig. 3**). In males, the WV-OD also induced glucose intolerance, though the increase in glucose excursion (area under the curve) was less pronounced than in the HFD group, while fasting blood glucose levels were similarly elevated (**Fig. 3 A**, **C**, **E**). In contrast, female mice fed the WV-OD maintained fasting blood glucose levels similar to controls and demonstrated only a non-significant trend toward impaired glucose tolerance, consistent with the sex-specific effect on body weight (**Fig. 3B**, **D**, **F**). Extending the dietary intervention to 14 weeks failed to induce glucose intolerance in WV-OD-fed females (**Fig. 4**). Furthermore, feeding the WV-OD to these mice did not alter serum lipids (**Fig. 5D-F**), organ weights (**Fig. 6B**), liver histology (**Fig. 6G, I**) or hepatic lipid content (**Fig. 6M-N**), in contrast to the effects observed with the HFD and consistent with their resistance to the WV-OD (**Fig. 2**).

**Figure 3.**
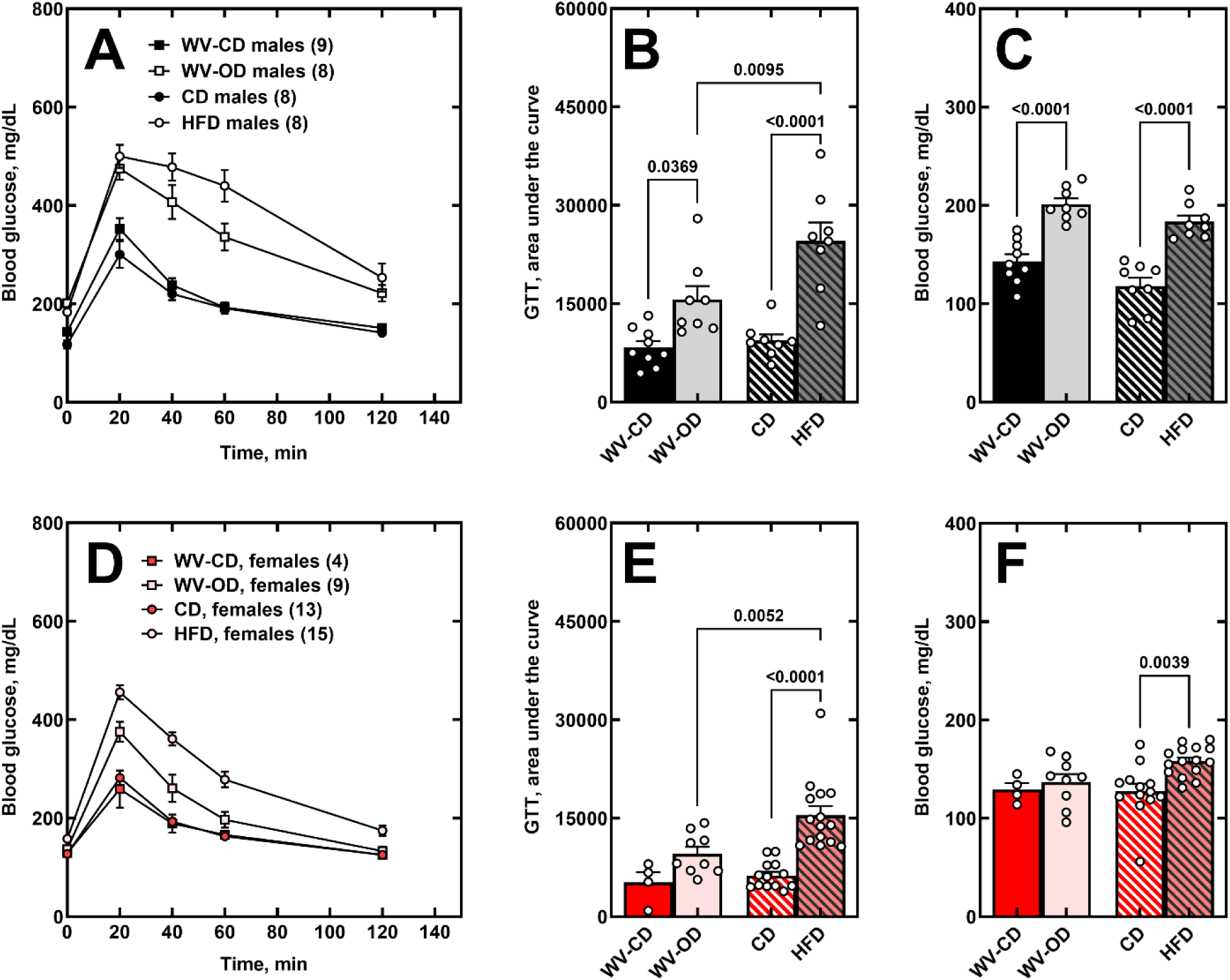
WV-OD induces glucose intolerance in male but not female mice after 10 weeks. (**A**, **D**) Glucose tolerance tests, (**B**, **E**) corresponding areas under the curve, and (**C**, **F**) fasting blood glucose levels in (**A-C**) male and (**D-F**) female mice after 10 weeks on the diets. Data are shown as the mean (bars) of individual animals (circles) ± SEM. (B, C, E, F) One-way ANOVA. p-values are shown for statistically significant comparisons.

**Figure 4.**
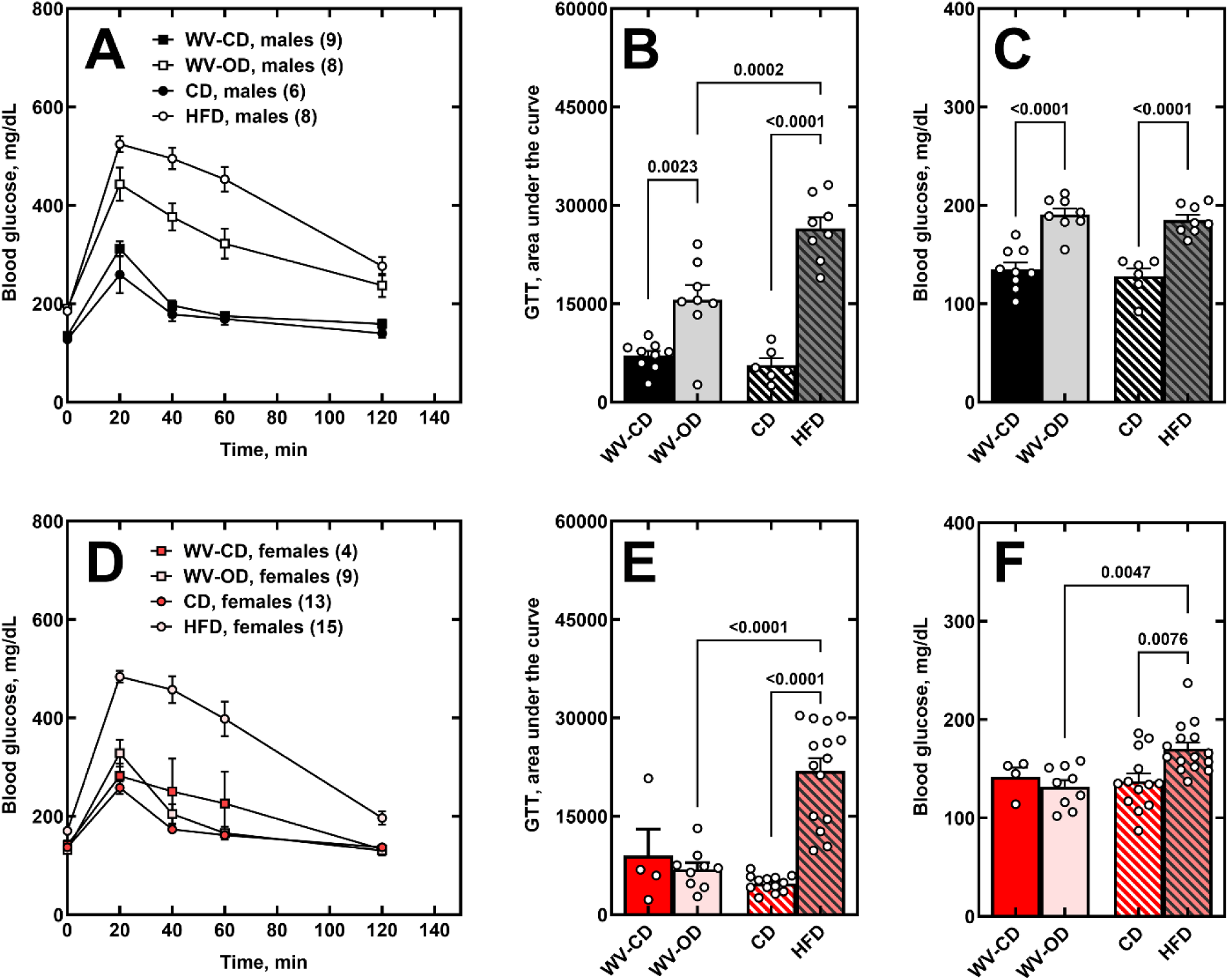
Extending WV-OD feeding to 14 weeks did not cause glucose intolerance in female mice. (**A**, **D**) Glucose tolerance tests, (**B**, **E**) corresponding areas under the curve, and (**C**, **F**) fasting blood glucose levels in (**A-C**) male and (**D-F**) female mice after 10 weeks on the diets. Data are shown as the mean (bars) of individual animals (circles) ± SEM. (B, C, E, F) One-way ANOVA. p-values are shown for statistically significant comparisons.

**Figure 5.**
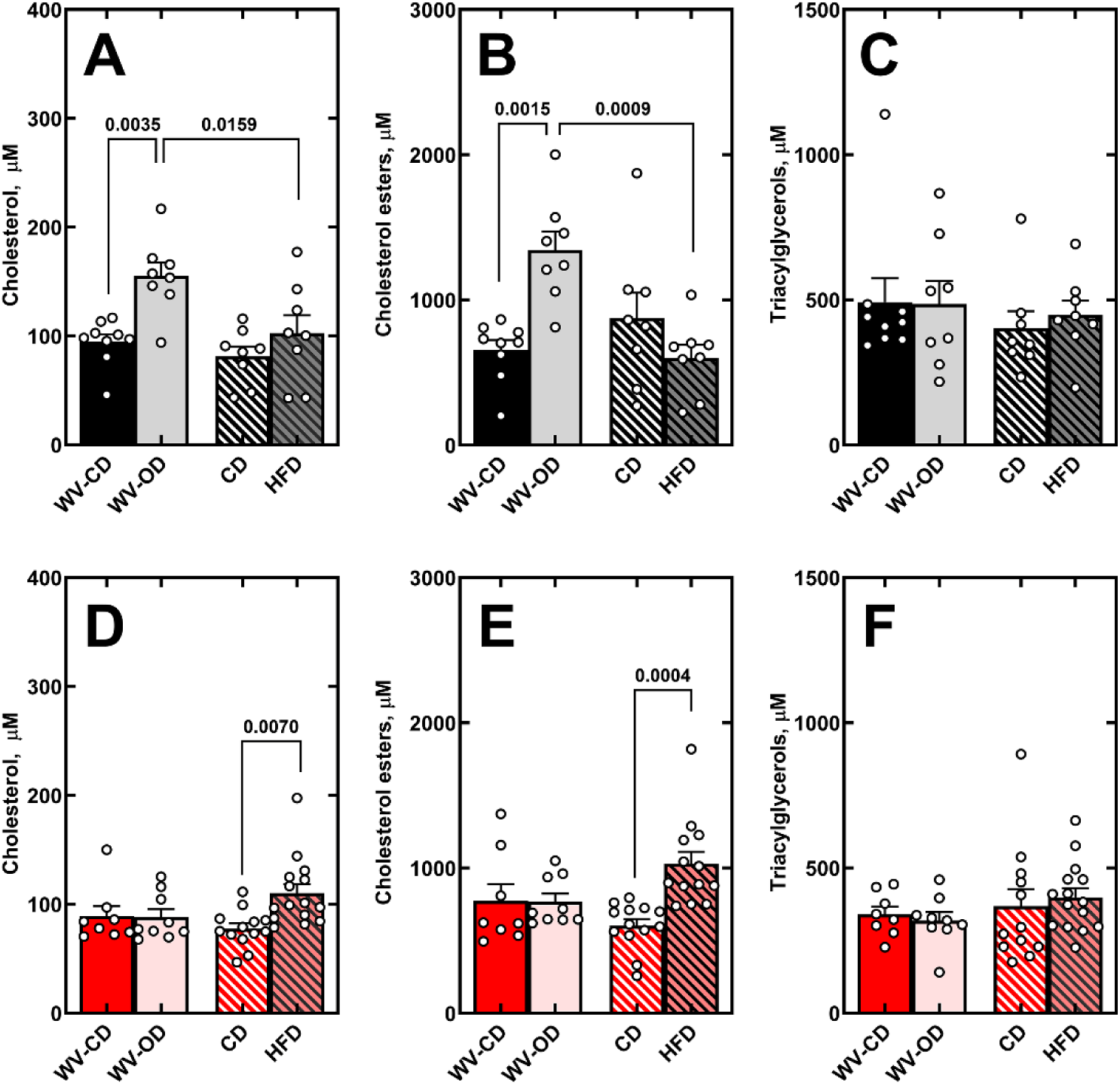
WV-OD increases circulating cholesterol and cholesterol esters in males. (**A**, **D**) Plasma cholesterol, (**B**, **E**) cholesterol esters, (**C**, **F**) and triglycerides in (**A-C**) male and (**D-F**) female mice. Data are shown as the mean (bars) of individual animals (circles) ± SEM. One-way ANOVA. p-values are shown for statistically significant comparisons.

**Figure 6.**
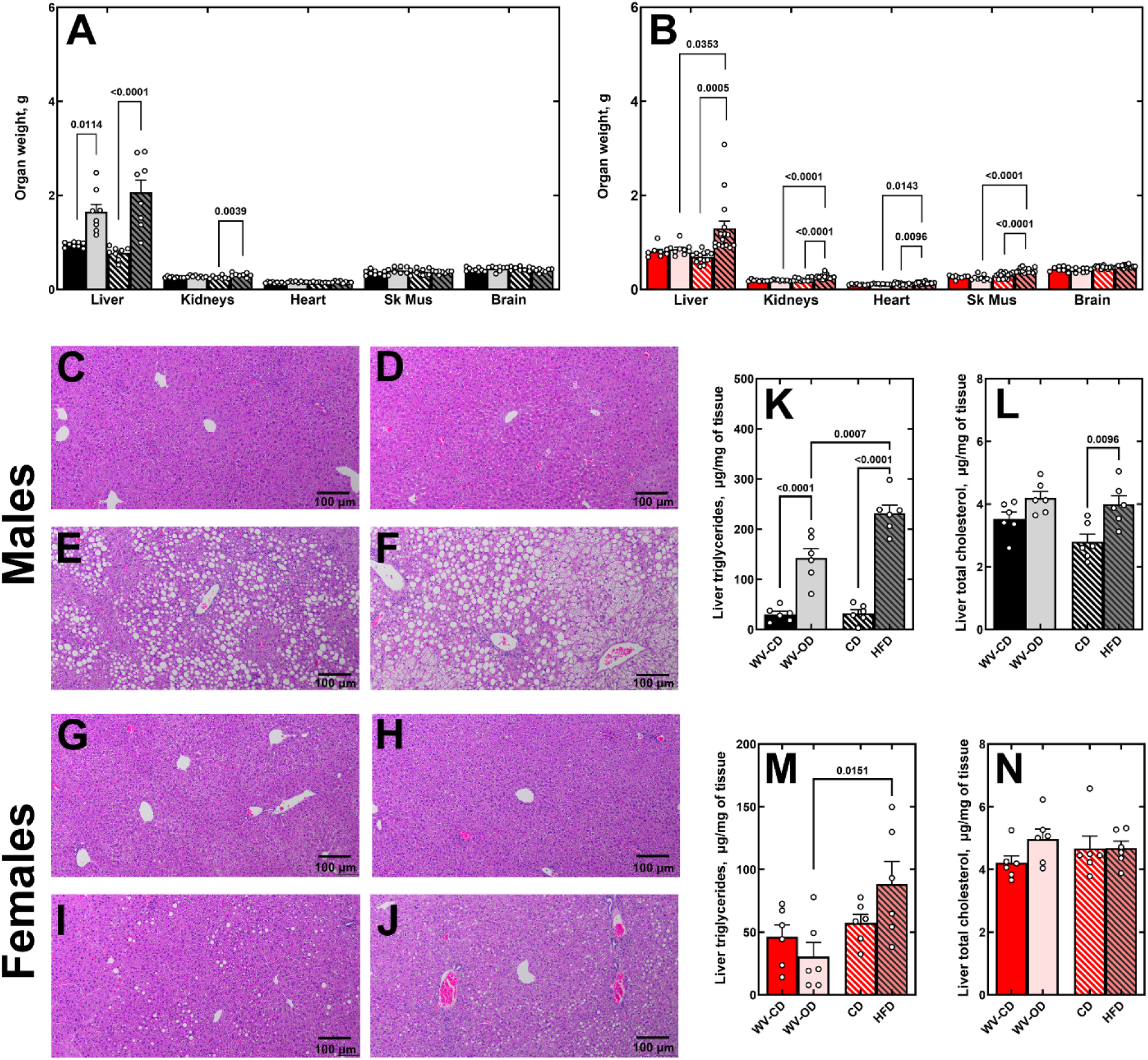
Effect of the WV-OD on organ weights and liver lipids. (**A**, **B**) Organ weights in (**A**) male and (**B**) female mice after 19 weeks of diet feeding. (**C–J**) Representative H&E-stained liver sections from (**C–F**) male and (**G–J**) female mice. Mice were fed (**C**, **G**) WV-CD, (**D**, **H**) CD, (**E**, **I**) WV-OD, and (**F**, **J**) HFD. (**K**, **M**) Liver triglycerides and (**L**, **N**) total cholesterol in (**K**, **L**) male and (**M**, **N**) female mice. Data are shown as the mean (bars) of individual animals (circles) ± SEM. One-way ANOVA. p-values are shown for statistically significant comparisons. Scale bars: 100 μm.

Male mice fed the WV-OD exhibited elevated circulating levels of cholesterol and cholesterol esters; an effect not observed in HFD-fed males (**Fig. 5A, B**)—despite no significant change in liver cholesterol content (**Fig. 6L**). Interestingly, although the HFD contains more dietary cholesterol than the WV-OD (0.028% vs. 0.020%), HFD-fed mice maintained normal serum cholesterol and cholesterol ester levels while accumulating significantly more cholesterol in the liver. This suggests that excess lipids in HFD-fed mice may be preferentially sequestered in the liver rather than circulating in the blood. Liver triglycerides were elevated in both diet groups, though the increase was more pronounced in HFD-fed mice (**Fig. 6K**). Consistent with the changes in adiposity, liver weights were significantly increased in both groups relative to their respective controls and were comparable between the WV-OD and HFD groups. Liver histology revealed consistent macrovesicular steatosis in male mice fed either high-fat diet, with no apparent difference in severity between the HFD and WV-OD groups, and markedly greater lipid accumulation compared to their respective control diets (**Fig. 6C-F**). In contrast, livers from female mice fed the HFD or WV-OD displayed only occasional microvesicular steatosis, with no notable differences between groups (**Fig. 6G-J**). Across all experimental conditions, there was no consistent evidence of diet-induced hepatic inflammation, regardless of sex or diet.

Combined, these results indicate that exposure to the WV-OD for 19 weeks leads to the development of glucose intolerance (10 and 14 weeks), accumulation of triglycerides in the liver, and elevated total cholesterol in plasma in male mice.

### WV-OD does not affect XOR activity or UA levels in liver and plasma

Diet-induced or genetic obesity models in rodents commonly exhibit increased liver and plasma UA levels, accompanied by upregulated XOR expression [29, 41–45]. This effect was reproduced in both male and female mice fed the HFD (**Fig. 7A, B, E, F**). Plasma UA levels were also significantly elevated in these mice, compared to controls (**Fig. 7D, H**), while circulating xanthine oxidase (XO) activity was selectively increased in HFD-fed females. In contrast, feeding the WV-OD did not alter hepatic XOR activity, plasma or hepatic UA levels, or circulating XO activity relative to WV-CD-fed controls. Notably, this lack of effect was also observed in WV-OD-fed male mice, despite having comparable body weight to HFD-fed mice and exhibiting robust hepatic triglyceride accumulation. These findings suggest that obesity alone is not sufficient to drive UA accumulation and XOR activation in this model.

**Figure 7.**
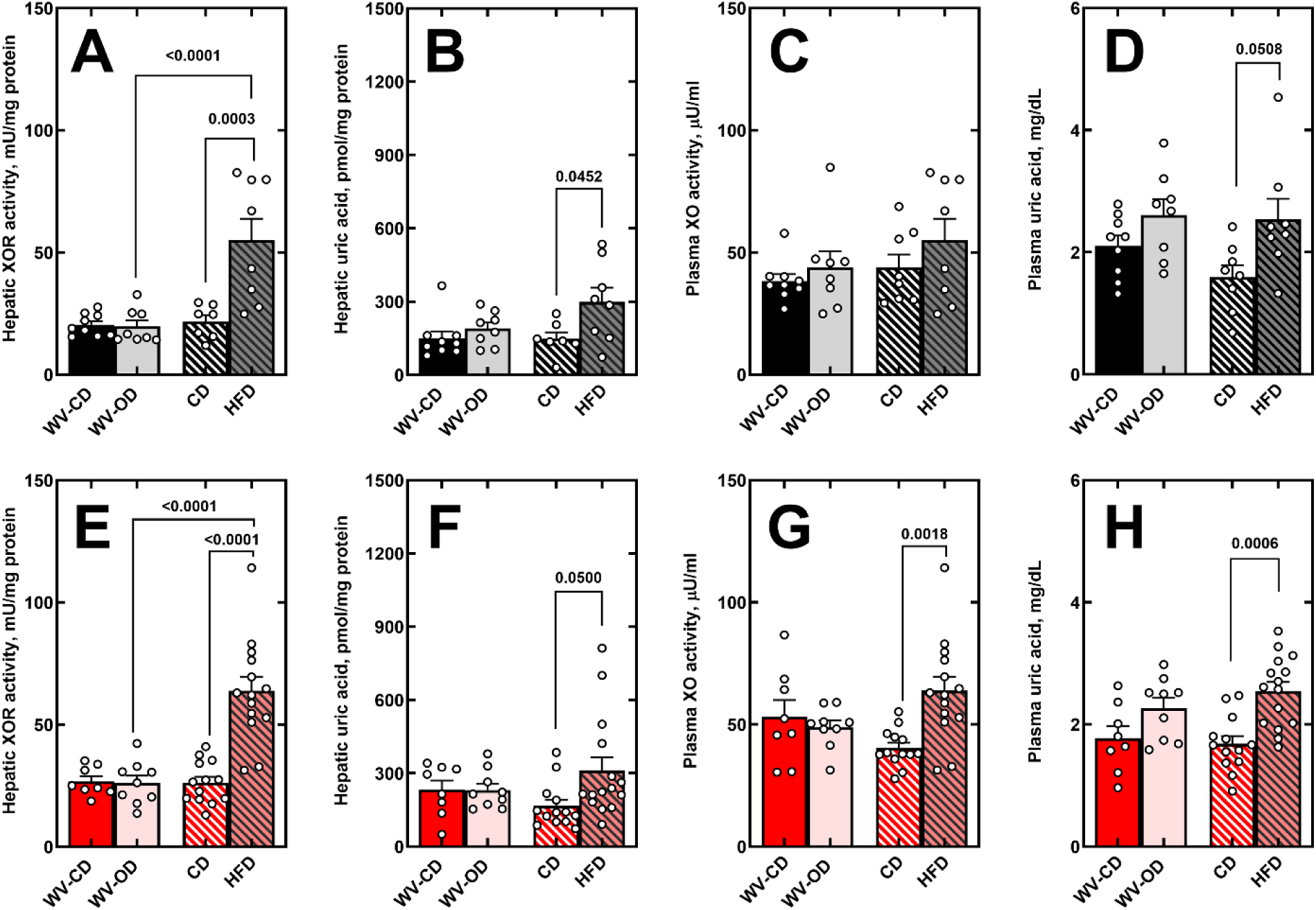
XOR activity and uric acid metabolism in liver and plasma. (**A**, **E**) Liver XOR activity and (**B**, **F**) uric acid levels in (**A**, **B**) male and (**E**, **F**) female mice. (**C**, **G**) Plasma XO activity and (**D**, **H**) uric acid levels in (**C**, **D**) male and (**G**, **H**) female mice. Data are shown as the mean (bars) of individual animals (circles) ± SEM. One-way ANOVA. p-values are shown for statistically significant comparisons.

## Discussion

In this study, we introduce the WV-OD, a regionally inspired, compositionally defined diet that models both the general features of Western dietary patterns and specific nutritional attributes reflective of high-risk populations. The WV-OD was formulated based on the nutritional analysis of meals reported by obese individuals in West Virginia, a state with the greatest prevalence (41.4%) of adult obesity and diabetes in the United States [21, 22]. While recent Western-style rodent diets such as the Total Western Diet (TWD), Standard American Diet (SAD), and modified SAD (mSAD) were developed using nationally aggregated survey data (e.g., NHANES) to reflect average U.S. macronutrient intake, only the SAD and mSAD have reliably induced metabolic dysfunction, likely due to the inclusion of elements typical of greater-risk subpopulations, such as trans-fats or high-fructose corn syrup [15, 16, 18, 19]. The WV-OD takes a distinct approach by basing its formulation on real-world meal data from a regional cohort, thereby capturing both the average macronutrient profile of the American diet and region-specific deviations such as lower fiber content and elevated sodium intake. To enable rigorous comparisons, we also developed a matched control diet (WV-CD) using the same ingredients and fat sources, thereby isolating the obesogenic effects attributable to nutrient quantity.

Our findings show that the WV-OD induces a robust, obese phenotype in male C57BL/6J mice, marked by increased body weight, adiposity, glucose intolerance, hepatic triglyceride accumulation, and elevated circulating cholesterol and cholesterol esters compared to WV-CD-fed controls. These effects were observed despite the WV-OD’s moderate fat content (39% kcal from fat), which is substantially less than the 60% kcal from fat in the commonly used HFD. Strikingly, male mice fed the WV-OD gained as much weight and demonstrated as much fat accretion as those fed the HFD, highlighting the obesogenic potency of a diet formulated to reflect real-world consumption patterns rather than extreme macronutrient loads. In contrast, female mice were largely resistant to the WV-OD when exposed for 19 weeks. Despite similar energy intake, only HFD-fed females exhibited significant weight gain and adiposity, suggesting that differences in nutrient absorption, energy expenditure, or other physiological factors may underlie the lack of phenotype in WV-OD-fed females. In addition, it is important to recognize that when compared to males, female mice require a greater time of exposure to high-fat feeding to demonstrate similar weight gain and metabolic dysfunction that males. This is evidenced by the final weights of males (45.3 ± 1.4 g) and females (40.0 ± 2.1 g) on the 60% HFD (**Fig. 2A&D**) as well as by previous reports demonstrating slower response curves for C57BL/6J females [28, 29].

Beyond total fat content, key compositional differences, such as fat type and the presence of soluble fiber, may have contributed to the sex disparity between males and females subjected to the WV-OD. Saturated fat has been more strongly associated with weight gain than unsaturated fat in both women and female rodents [46, 47]. The WV-OD contains significantly less saturated fat (20% of total fat by weight, 40% saturated) (**Fig. 1C**) than the HFD (35% of total fat by weight, primarily saturated). Additionally, emerging evidence identifies the gut microbiota as a critical determinant of weight gain resistance in females [48, 49]. The inclusion of inulin in the WV-OD may have contributed to this protection by promoting a healthier gut microbiome. Moreover, inulin has been shown to prevent intestinal atrophy and metabolic disturbances associated with cellulose-only purified diets like the HFD [38, 50]. Future studies will examine gut microbial composition and microbiota-derived metabolites in WV-OD versus HFD-fed mice to test this hypothesis directly. Overall, the resistance of female mice to developing obesity when fed the WV-OD is at odds with obesity rates reported among female residents of West Virginia and underscores the importance of identifying mechanisms that limit the translatability of female murine models, which could also include housing at temperatures below the thermoneutral zone (∼30 °C for mice), particularly given known sex differences in thermogenesis [51–53].

Even in the context of comparable weight gain in male mice, compositional differences between the WV-OD and the HFD produced distinct metabolic signatures. Notably, only the WV-OD elevated circulating cholesterol and cholesterol esters, despite containing less dietary cholesterol. These measurements were taken after a 7-hour fast, minimizing chylomicron contribution and suggesting that the differences reflect changes in lipoprotein metabolism rather than postprandial lipid absorption and transport. However, because lipoprotein fractionation was not performed, we cannot say whether these increases were driven by specific lipoprotein classes such as HDL or LDL. Conversely, only HFD-fed mice exhibited increased xanthine oxidoreductase (XOR) activity and elevated UA levels in the liver, effects absent in WV-OD-fed animals despite similar levels of obesity and hepatic lipid accumulation. A similar pattern was seen for XOR and UA in the circulation; however, the WV-OD (males and females), while statistically not different from their controls, did demonstrate a trend towards elevation in plasma UA. This is an important point as rodent models of diet-induced obesity as well as models of genetic-induced obesity (*e.g.* ob/ob, db/db and pound mouse) have demonstrated obesity to associate tightly with elevation in circulating UA levels [29, 41–45]. These observations, combined with the realization that human obesity and diabetes also demonstrate elevation of plasma UA [54–57], have identified UA as a biomarker of obesity and incentivized the hypothesis that UA may be causative of the metabolic consequences allied to obesity [58, 59]. While it has been reported that interventions to maintain lean levels of circulating UA in the context of obesity do not impact metabolic dysfunction [42], further investigation is needed before a consensus can be reached on whether the relationship between UA and obesity is causal or merely correlative.

A few limitations of this study should be acknowledged. *First*, the WV-OD was derived from the nutritional analysis of 27 meals, which represents a relatively small sample size and relies on dietary self-reporting, a process that inherently introduces recall bias and uncertainty in estimating true nutrient intake. While assumptions regarding portion size and food composition were applied consistently, inaccuracies in food reporting are difficult to avoid. The WV-OD was developed using region-specific dietary data and the results should be interpreted within the appropriate geographic context. Nevertheless, this approach underscores the value of regionally informed dietary models in capturing metabolic risk factors that may be underrepresented in nationally averaged diets. *Second*, by incorporating sucrose derived from SSBs directly into the solid diet, we may have attenuated some of the deleterious metabolic effects typically associated with liquid sucrose consumption [60]. A growing body of evidence demonstrates that sucrose ingested in liquid form promotes greater adiposity and hepatic steatosis than equivalent amounts delivered in solid food, likely due to differences in satiety, caloric compensation, and hepatic carbohydrate flux [61–63]. From this perspective, the WV-OD represents a conservative yet physiologically relevant model of dietary sugar exposure.

*In toto*, the WV-OD is a newly developed diet that captures both the composition and metabolic consequences of dietary patterns observed in overweight and obese individuals that is specific to local dietary habits. By balancing regional specificity with broader national relevance, the WV-OD provides a valuable preclinical model for diet-induced obesity and metabolic dysfunction. It also serves to further amplify the impact of sex on the development of obesity in preclinical models and provides fertile ground for further investigation designed to unravel these sex-related differences.

## Abbreviations

GTT: glucose tolerance test
HFD: high-fat diet
WV-OD: West Virginia obesogenic diet
UA: uric acid
XDH: xanthine dehydrogenase
XO: xanthine oxidase
XOR: xanthine oxidoreductase

## Author contributions

Concept: RL, EEK, JMH, PM; Design: RL, EEK, APG; Data collection, analysis, and interpretation: EEK, APG, BAM, ALS, SEL, SRS, MF, SB, PF, EAK, DCS, FJS, NKHK, RL; Writing: EEK, APG, RL.

## Data availability statement

All data supporting the findings of this study are included in the manuscript and supplementary materials, either as summary graphs or in tabular data files.

## Supplemental Figures

**Supplemental Figure S1.**
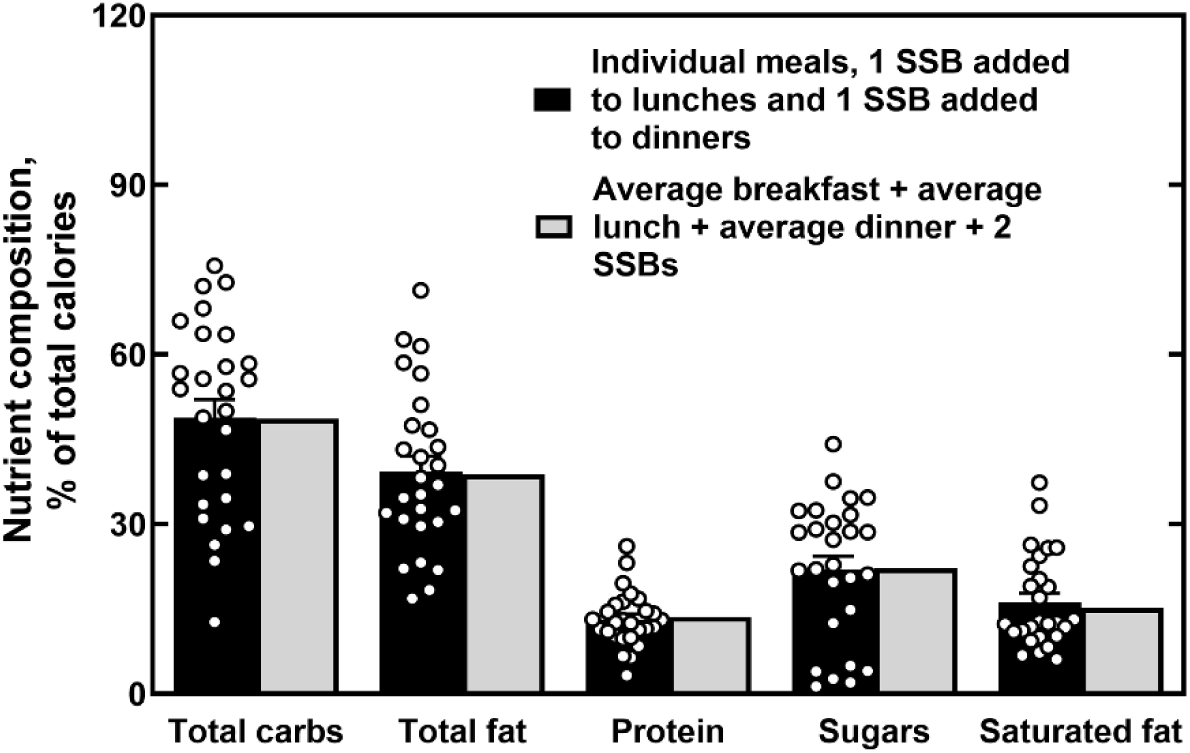
Compositional analysis of meals using two methods for incorporating sugar-sweetened beverages (SSBs). Caloric contributions from total carbohydrates, sugars, total fat, saturated fat, and protein were estimated using two approaches for incorporating SSBs. Method 1: one SSB was added to each lunch and dinner before calculating the caloric contribution of each nutrient for every individual meal analyzed (circles). Method 2: two SSBs were added to the average nutrient composition of breakfast, lunch, and dinner meals.

**Supplemental Figure S2.**
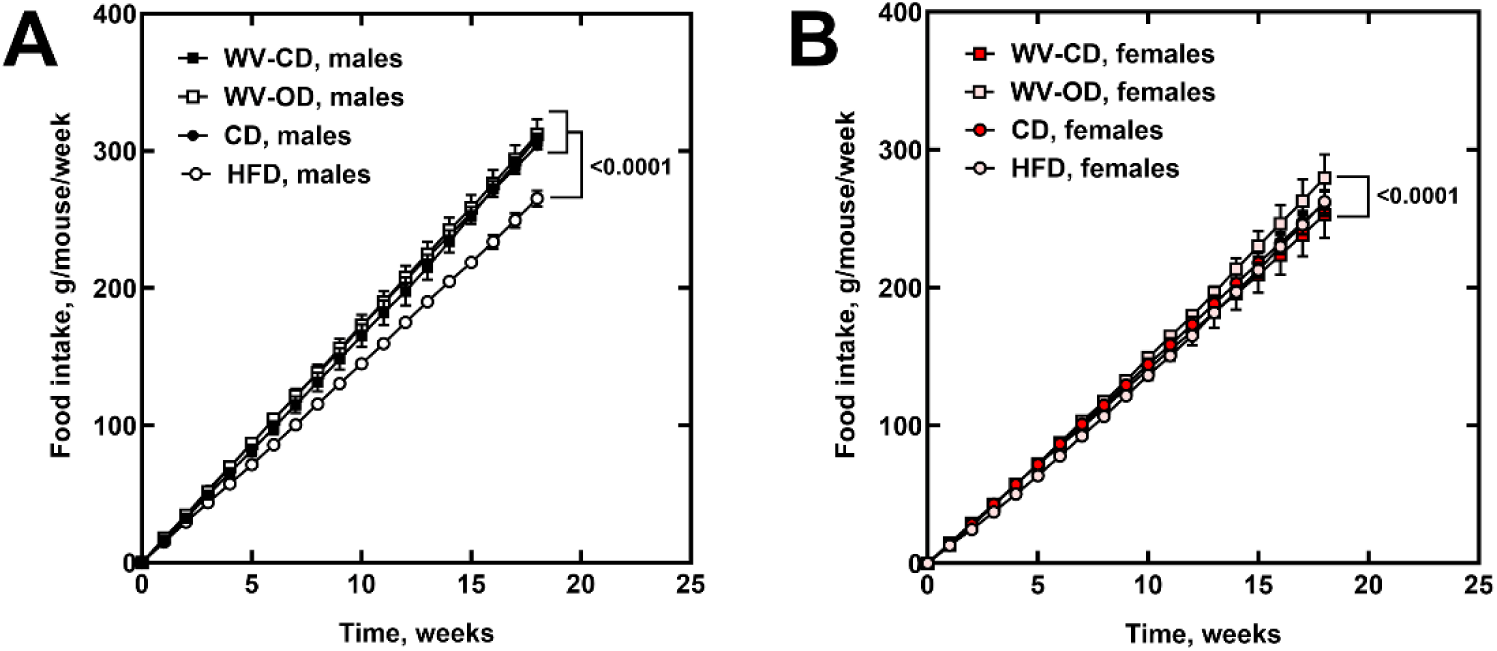
Cumulative food intake over the study period. Food consumption over 19 weeks of diet feeding in (**A**) male and (**B**) female mice. Linear regression with slope comparison; p-values are shown for statistically significant comparisons.

## References

1. The Lancet Gastroenterology H. Obesity: another ongoing pandemic. Lancet Gastroenterol Hepatol. 2021;6(6):411. doi: 10.1016/S2468-1253(21)00143-6. PubMed PMID: 34015350; PubMed Central PMCID: PMCPMC9259282.

2. Collaboration NCDRF. Worldwide trends in underweight and obesity from 1990 to 2022: a pooled analysis of 3663 population-representative studies with 222 million children, adolescents, and adults. Lancet. 2024;403(10431):1027–50. Epub 20240229. doi: 10.1016/S0140-6736(23)02750-2. PubMed PMID: 38432237; PubMed Central PMCID: PMCPMC7615769.

3. Emmerich SD, Fryar, C.D., Stierman, B., Ogden, C.L. Obesity and Severe Obesity Prevalence in Adults: United States, August 2021–August 2023. NCHS Data Brief no 508. 2024. doi: 10.15620/cdc/159281.

4. Fryar CD, Carroll, M.D., Afful, J. Prevalence of Overweight, Obesity, and Severe Obesity Among Adults Aged 20 and Over: United States, 1960–1962 Through 2017–2018. NCHS Health E-Stats. 2020.

5. Collaborators GUOF. National-level and state-level prevalence of overweight and obesity among children, adolescents, and adults in the USA, 1990-2021, and forecasts up to 2050. Lancet. 2024;404(10469):2278–98. Epub 20241114. doi: 10.1016/S0140-6736(24)01548-4. PubMed PMID: 39551059; PubMed Central PMCID: PMCPMC11694015.

6. Buettner R, Scholmerich J, Bollheimer LC. High-fat diets: modeling the metabolic disorders of human obesity in rodents. Obesity (Silver Spring). 2007;15(4):798–808. doi: 10.1038/oby.2007.608. PubMed PMID: 17426312.

7. Hariri N, Thibault L. High-fat diet-induced obesity in animal models. Nutr Res Rev. 2010;23(2):270–99. Epub 20101027. doi: 10.1017/S0954422410000168. PubMed PMID: 20977819.

8. Speakman JR. Use of high-fat diets to study rodent obesity as a model of human obesity. Int J Obes (Lond). 2019;43(8):1491–2. Epub 20190409. doi: 10.1038/s41366-019-0363-7. PubMed PMID: 30967607.

9. Johnston SL, Souter DM, Tolkamp BJ, Gordon IJ, Illius AW, Kyriazakis I, Speakman JR. Intake compensates for resting metabolic rate variation in female C57BL/6J mice fed high-fat diets. Obesity (Silver Spring). 2007;15(3):600–6. doi: 10.1038/oby.2007.550. PubMed PMID: 17372309.

10. Ha SK, Chae C. Inducible nitric oxide distribution in the fatty liver of a mouse with high fat diet-induced obesity. Exp Anim. 2010;59(5):595–604. doi: 10.1538/expanim.59.595. PubMed PMID: 21030787.

11. Skalski HJ, Arendt AR, Harkins SK, MacLachlan M, Corbett CJM, Goy RW, et al. Key Considerations for Studying the Effects of High-Fat Diet on the Nulligravid Mouse Endometrium. J Endocr Soc. 2024;8(7):bvae104. Epub 20240525. doi: 10.1210/jendso/bvae104. PubMed PMID: 38854907; PubMed Central PMCID: PMCPMC11156617.

12. Sclafani A, Springer D. Dietary obesity in adult rats: similarities to hypothalamic and human obesity syndromes. Physiol Behav. 1976;17(3):461–71. doi: 10.1016/0031-9384(76)90109-8. PubMed PMID: 1013192.

13. Lalanza JF, Snoeren EMS. The cafeteria diet: A standardized protocol and its effects on behavior. Neurosci Biobehav Rev. 2021;122:92–119. Epub 20201210. doi: 10.1016/j.neubiorev.2020.11.003. PubMed PMID: 33309818.

14. Sampey BP, Vanhoose AM, Winfield HM, Freemerman AJ, Muehlbauer MJ, Fueger PT, et al. Cafeteria diet is a robust model of human metabolic syndrome with liver and adipose inflammation: comparison to high-fat diet. Obesity (Silver Spring). 2011;19(6):1109–17. Epub 20110217. doi: 10.1038/oby.2011.18. PubMed PMID: 21331068; PubMed Central PMCID: PMCPMC3130193.

15. Hintze KJ, Benninghoff AD, Ward RE. Formulation of the Total Western Diet (TWD) as a basal diet for rodent cancer studies. J Agric Food Chem. 2012;60(27):6736–42. Epub 20120124. doi: 10.1021/jf204509a. PubMed PMID: 22224871.

16. Monsanto SP, Hintze KJ, Ward RE, Larson DP, Lefevre M, Benninghoff AD. The new total Western diet for rodents does not induce an overweight phenotype or alter parameters of metabolic syndrome in mice. Nutr Res. 2016;36(9):1031–44. Epub 20160606. doi: 10.1016/j.nutres.2016.06.002. PubMed PMID: 27632924.

17. Totsch SK, Quinn TL, Strath LJ, McMeekin LJ, Cowell RM, Gower BA, Sorge RE. The impact of the Standard American Diet in rats: Effects on behavior, physiology and recovery from inflammatory injury. Scand J Pain. 2017;17:316–24. Epub 20170918. doi: 10.1016/j.sjpain.2017.08.009. PubMed PMID: 28927908.

18. Totsch SK, Meir RY, Quinn TL, Lopez SA, Gower BA, Sorge RE. Effects of a Standard American Diet and an anti-inflammatory diet in male and female mice. Eur J Pain. 2018;22(7):1203–13. Epub 20180302. doi: 10.1002/ejp.1207. PubMed PMID: 29436058.

19. Chehade SB, Green GBH, Graham CD, Chakraborti A, Vashai B, Moon A, et al. A modified standard American diet induces physiological parameters associated with metabolic syndrome in C57BL/6J mice. Front Nutr. 2022;9:929446. Epub 20220829. doi: 10.3389/fnut.2022.929446. PubMed PMID: 36105576; PubMed Central PMCID: PMCPMC9464921.

20. Reeves PG. Components of the AIN-93 diets as improvements in the AIN-76A diet. J Nutr. 1997;127(5 Suppl):838S-41S. doi: 10.1093/jn/127.5.838S. PubMed PMID: 9164249.

21. Warren MSM, West, M. The State of Obesity 2025: Better Policies for a Healthier America. 2025. doi: https://www.tfah.org/.

22. Incidence of Diagnosed Diabetes in Adults, Aged 18-76 Years Old, West Virginia, 2021 2021. Available from: https://nccd.cdc.gov/Toolkit/DiabetesBurden/Incidence/.

23. Lundeen EA, Park S, Pan L, Blanck HM. Daily Intake of Sugar-Sweetened Beverages Among US Adults in 9 States, by State and Sociodemographic and Behavioral Characteristics, 2016. Prev Chronic Dis. 2018;15:E154. doi: 10.5888/pcd15.180335. PubMed PMID: 30576280; PubMed Central PMCID: PMCPMC6307838.

24. Dietary Guidelines for Americans, 2020-2025. US Department of Agriculture and US Department of Health and Human Services. 2020;9th Edition.

25. Nutrient Requirements of Laboratory Animals: Fourth Revised Edition, 1995. Washington (DC)1995.

26. Vickers SD, Shumar SA, Saporito DC, Kunovac A, Hathaway QA, Mintmier B, et al. NUDT7 regulates total hepatic CoA levels and the composition of the intestinal bile acid pool in male mice fed a Western diet. J Biol Chem. 2023;299(1):102745. doi: 10.1016/j.jbc.2022.102745. PubMed PMID: 36436558; PubMed Central PMCID: PMCPMC9792899.

27. Lewis SE, Rosencrance CB, De Vallance E, Giromini A, Williams XM, Oh JY, et al. Human and rodent red blood cells do not demonstrate xanthine oxidase activity or XO-catalyzed nitrite reduction to NO. Free Radic Biol Med. 2021;174:84–8. Epub 20210715. doi: 10.1016/j.freeradbiomed.2021.07.012. PubMed PMID: 34273539; PubMed Central PMCID: PMCPMC9257433.

28. Lewis SE, Li L, Fazzari M, Salvatore SR, Li J, Hileman EA, et al. Obese female mice do not exhibit overt hyperuricemia despite hepatic steatosis and impaired glucose tolerance. Adv Redox Res. 2022;6. Epub 20221007. doi: 10.1016/j.arres.2022.100051. PubMed PMID: 36561324; PubMed Central PMCID: PMCPMC9770588.

29. Giromini AP, Salvatore, S.R., Maxwell, B.A., Lewis, S.E., Gunther, M.R., Fazzari, M., Schopfer, F.J., Leonardi, R., Kelley, E.E. Obesity-Associated Hyperuricemia in Female Mice: A Reevaluation. Gout, Urate, and Crystal Deposition Disease. 2024;2(3):252–65. doi: 10.3390/gucdd2030019 -.

30. Shumar SA, Kerr EW, Fagone P, Infante AM, Leonardi R. Overexpression of Nudt7 decreases bile acid levels and peroxisomal fatty acid oxidation in the liver. J Lipid Res. 2019;60(5):1005–19. doi: 10.1194/jlr.M092676. PubMed PMID: 30846528; PubMed Central PMCID: PMCPMC6495166.

31. Lepage G, Roy CC. Direct transesterification of all classes of lipids in a one-step reaction. J Lipid Res. 1986;27(1):114–20. PubMed PMID: 3958609.

32. Tran QD, Nguyen THH, Le CL, Hoang LV, Vu TQC, Phan NQ, Bui TT. Sugar-sweetened beverages consumption increases the risk of metabolic syndrome and its components in adults: Consistent and robust evidence from an umbrella review. Clin Nutr ESPEN. 2023;57:655–64. Epub 20230812. doi: 10.1016/j.clnesp.2023.08.001. PubMed PMID: 37739720.

33. Nguyen M, Jarvis SE, Tinajero MG, Yu J, Chiavaroli L, Mejia SB, et al. Sugar-sweetened beverage consumption and weight gain in children and adults: a systematic review and meta-analysis of prospective cohort studies and randomized controlled trials. Am J Clin Nutr. 2023;117(1):160–74. Epub 20221220. doi: 10.1016/j.ajcnut.2022.11.008. PubMed PMID: 36789935.

34. Santos LP, Gigante DP, Delpino FM, Maciel AP, Bielemann RM. Sugar sweetened beverages intake and risk of obesity and cardiometabolic diseases in longitudinal studies: A systematic review and meta-analysis with 1.5 million individuals. Clin Nutr ESPEN. 2022;51:128–42. Epub 20220824. doi: 10.1016/j.clnesp.2022.08.021. PubMed PMID: 36184197.

35. Chevinsky JR, Lee SH, Blanck HM, Park S. Prevalence of Self-Reported Intake of Sugar-Sweetened Beverages Among US Adults in 50 States and the District of Columbia, 2010 and 2015. Prev Chronic Dis. 2021;18:E35. doi: 10.5888/pcd18.200434. PubMed PMID: 33856977; PubMed Central PMCID: PMCPMC8051857.

36. Simopoulos AP. The importance of the ratio of omega-6/omega-3 essential fatty acids. Biomed Pharmacother. 2002;56(8):365–79. doi: 10.1016/s0753-3322(02)00253-6. PubMed PMID: 12442909.

37. Crick DCP, Halligan SL, Davey Smith G, Khandaker GM, Jones HJ. The relationship between polyunsaturated fatty acids and inflammation: evidence from cohort and Mendelian randomization analyses. Int J Epidemiol. 2025;54(4). doi: 10.1093/ije/dyaf065. PubMed PMID: 40550517; PubMed Central PMCID: PMCPMC12204398.

38. Chassaing B, Miles-Brown J, Pellizzon M, Ulman E, Ricci M, Zhang L, et al. Lack of soluble fiber drives diet-induced adiposity in mice. Am J Physiol Gastrointest Liver Physiol. 2015;309(7):G528-41. Epub 20150716. doi: 10.1152/ajpgi.00172.2015. PubMed PMID: 26185332; PubMed Central PMCID: PMCPMC4593822.

39. Juskiewicz J, Zdunczyk Z. Effects of cellulose, carboxymethylcellulose and inulin fed to rats as single supplements or in combinations on their caecal parameters. Comp Biochem Physiol A Mol Integr Physiol. 2004;139(4):513–9. doi: 10.1016/j.cbpb.2004.10.015. PubMed PMID: 15596397.

40. Hu S, Wang L, Yang D, Li L, Togo J, Wu Y, et al. Dietary Fat, but Not Protein or Carbohydrate, Regulates Energy Intake and Causes Adiposity in Mice. Cell Metab. 2018;28(3):415–31 e4. Epub 20180712. doi: 10.1016/j.cmet.2018.06.010. PubMed PMID: 30017356.

41. Khavjou O, Tayebali Z, Cho P, Myers K, Zhang P. Rural-Urban Disparities in State-Level Diabetes Prevalence Among US Adults, 2021. Prev Chronic Dis. 2025;22:E05. Epub 20250116. doi: 10.5888/pcd22.240199. PubMed PMID: 39819894; PubMed Central PMCID: PMCPMC11870018.

42. Harmon DB, Mandler WK, Sipula IJ, Dedousis N, Lewis SE, Eckels JT, et al. Hepatocyte-Specific Ablation or Whole-Body Inhibition of Xanthine Oxidoreductase in Mice Corrects Obesity-Induced Systemic Hyperuricemia Without Improving Metabolic Abnormalities. Diabetes. 2019;68(6):1221–9. Epub 20190401. doi: 10.2337/db18-1198. PubMed PMID: 30936145; PubMed Central PMCID: PMCPMC6610025.

43. Battelli MG, Bortolotti M, Polito L, Bolognesi A. The role of xanthine oxidoreductase and uric acid in metabolic syndrome. Biochim Biophys Acta Mol Basis Dis. 2018;1864(8):2557–65. Epub 20180505. doi: 10.1016/j.bbadis.2018.05.003. PubMed PMID: 29733945.

44. Kelley EE, Baust J, Bonacci G, Golin-Bisello F, Devlin JE, St Croix CM, et al. Fatty acid nitroalkenes ameliorate glucose intolerance and pulmonary hypertension in high-fat diet-induced obesity. Cardiovasc Res. 2014;101(3):352–63. Epub 20140102. doi: 10.1093/cvr/cvt341. PubMed PMID: 24385344; PubMed Central PMCID: PMCPMC3928004.

45. Lanaspa MA, Sanchez-Lozada LG, Choi YJ, Cicerchi C, Kanbay M, Roncal-Jimenez CA, et al. Uric acid induces hepatic steatosis by generation of mitochondrial oxidative stress: potential role in fructose-dependent and -independent fatty liver. J Biol Chem. 2012;287(48):40732–44. Epub 20121003. doi: 10.1074/jbc.M112.399899. PubMed PMID: 23035112; PubMed Central PMCID: PMCPMC3504786.

46. Hariri N, Gougeon R, Thibault L. A highly saturated fat-rich diet is more obesogenic than diets with lower saturated fat content. Nutr Res. 2010;30(9):632–43. doi: 10.1016/j.nutres.2010.09.003. PubMed PMID: 20934605.

47. Field AE, Willett WC, Lissner L, Colditz GA. Dietary fat and weight gain among women in the Nurses’ Health Study. Obesity (Silver Spring). 2007;15(4):967–76. doi: 10.1038/oby.2007.616. PubMed PMID: 17426332.

48. Peng C, Xu X, Li Y, Li X, Yang X, Chen H, et al. Sex-specific association between the gut microbiome and high-fat diet-induced metabolic disorders in mice. Biol Sex Differ. 2020;11(1):5. Epub 20200120. doi: 10.1186/s13293-020-0281-3. PubMed PMID: 31959230; PubMed Central PMCID: PMCPMC6971877.

49. Czarnowski P, Balabas A, Kulaga Z, Kulecka M, Goryca K, Pysniak K, et al. Effects of Soluble Dextrin Fiber from Potato Starch on Body Weight and Associated Gut Dysbiosis Are Evident in Western Diet-Fed Mice but Not in Overweight/Obese Children. Nutrients. 2024;16(7). Epub 20240322. doi: 10.3390/nu16070917. PubMed PMID: 38612951; PubMed Central PMCID: PMCPMC11013109.

50. Schipke J, Brandenberger C, Vital M, Muhlfeld C. Starch and Fiber Contents of Purified Control Diets Differentially Affect Hepatic Lipid Homeostasis and Gut Microbiota Composition. Front Nutr. 2022;9:915082. Epub 20220707. doi: 10.3389/fnut.2022.915082. PubMed PMID: 35873446; PubMed Central PMCID: PMCPMC9301012.

51. Fernandez-Pena C, Reimundez A, Viana F, Arce VM, Senaris R. Sex differences in thermoregulation in mammals: Implications for energy homeostasis. Front Endocrinol (Lausanne). 2023;14:1093376. Epub 20230308. doi: 10.3389/fendo.2023.1093376. PubMed PMID: 36967809; PubMed Central PMCID: PMCPMC10030879.

52. Gomez-Garcia I, Trepiana J, Fernandez-Quintela A, Giralt M, Portillo MP. Sexual Dimorphism in Brown Adipose Tissue Activation and White Adipose Tissue Browning. Int J Mol Sci. 2022;23(15). Epub 20220726. doi: 10.3390/ijms23158250. PubMed PMID: 35897816; PubMed Central PMCID: PMCPMC9368277.

53. Ganeshan K, Chawla A. Warming the mouse to model human diseases. Nat Rev Endocrinol. 2017;13(8):458–65. doi: 10.1038/nrendo.2017.48. PubMed PMID: 28497813; PubMed Central PMCID: PMCPMC5777302.

54. Ebrahimpour P, Fakhrzadeh H, Heshmat R, Bandarian F, Larijani B. Serum uric acid levels and risk of metabolic syndrome in healthy adults. Endocr Pract. 2008;14(3):298–304. doi: 10.4158/EP.14.3.298. PubMed PMID: 18463036.

55. Tam HK, Kelly AS, Fox CK, Nathan BM, Johnson LA. Weight Loss Mediated Reduction in Xanthine Oxidase Activity and Uric Acid Clearance in Adolescents with Severe Obesity. Child Obes. 2016;12(4):286–91. Epub 20160315. doi: 10.1089/chi.2015.0051. PubMed PMID: 26978590; PubMed Central PMCID: PMCPMC5911696.

56. Li F, Chen S, Qiu X, Wu J, Tan M, Wang M. Serum Uric Acid Levels and Metabolic Indices in an Obese Population: A Cross-Sectional Study. Diabetes Metab Syndr Obes. 2021;14:627–35. Epub 20210211. doi: 10.2147/DMSO.S286299. PubMed PMID: 33603427; PubMed Central PMCID: PMCPMC7886379.

57. Li S, Feng L, Sun X, Ding J, Zhou W. Association between serum uric acid and measures of adiposity in Chinese adults: a cross-sectional study. BMJ Open. 2023;13(5):e072317. Epub 20230524. doi: 10.1136/bmjopen-2023-072317. PubMed PMID: 37225271; PubMed Central PMCID: PMCPMC10230993.

58. Nakagawa T, Hu H, Zharikov S, Tuttle KR, Short RA, Glushakova O, et al. A causal role for uric acid in fructose-induced metabolic syndrome. Am J Physiol Renal Physiol. 2006;290(3):F625–31. Epub 20051018. doi: 10.1152/ajprenal.00140.2005. PubMed PMID: 16234313.

59. Sanchez-Lozada LG, Tapia E, Bautista-Garcia P, Soto V, Avila-Casado C, Vega-Campos IP, et al. Effects of febuxostat on metabolic and renal alterations in rats with fructose-induced metabolic syndrome. Am J Physiol Renal Physiol. 2008;294(4):F710–8. Epub 20080123. doi: 10.1152/ajprenal.00454.2007. PubMed PMID: 18216151.

60. Kawasaki T, Kashiwabara A, Sakai T, Igarashi K, Ogata N, Watanabe H, et al. Long-term sucrose-drinking causes increased body weight and glucose intolerance in normal male rats. Br J Nutr. 2005;93(5):613–8. doi: 10.1079/bjn20051407. PubMed PMID: 15975159.

61. DiMeglio DP, Mattes RD. Liquid versus solid carbohydrate: effects on food intake and body weight. Int J Obes Relat Metab Disord. 2000;24(6):794–800. doi: 10.1038/sj.ijo.0801229. PubMed PMID: 10878689.

62. Malik VS, Hu FB. Fructose and Cardiometabolic Health: What the Evidence From Sugar-Sweetened Beverages Tells Us. J Am Coll Cardiol. 2015;66(14):1615–24. doi: 10.1016/j.jacc.2015.08.025. PubMed PMID: 26429086; PubMed Central PMCID: PMCPMC4592517.

63. Togo J, Hu S, Li M, Niu C, Speakman JR. Impact of dietary sucrose on adiposity and glucose homeostasis in C57BL/6J mice depends on mode of ingestion: liquid or solid. Mol Metab. 2019;27:22–32. Epub 20190604. doi: 10.1016/j.molmet.2019.05.010. PubMed PMID: 31255519; PubMed Central PMCID: PMCPMC6717800.

